# Recently evolved, stage-specific genes are enriched at life stage transitions in flies

**DOI:** 10.1101/2025.04.28.650992

**Authors:** Andreas Remmel, Karl K. Käther, Peter F. Stadler, Steffen Lemke

## Abstract

Understanding how genomic information is selectively utilized across different life stages is essential for deciphering the developmental and evolutionary strategies of metazoans. In holometabolous insects, the dynamic expression of genes enables distinct functional adaptations at embryonic, larval, pupal and adult stages, likely contributing to their evolutionary success. While *Drosophila melanogaster* has been extensively studied, less is known about the evolutionary dynamics that could govern stage-specific gene expression. To address this question, we compared the distribution of stage-specific genes, i.e., genes expressed in temporally restricted developmental stages, across development of *D.melanogaster* and *Aedes aegypti*. Using tau-scoring, a computational method to determine gene expression specificity, we found that, on average, a large proportion of genes (20 to 30% of all protein-coding genes) in both species exhibit restricted expression to specific developmental stages. Phylostratigraphy analysis, a method to date the age of genes, further revealed that stage-specific genes fall into two major categories: highly conserved and recently evolved. Notably, many of the recently evolved and stage-specific genes identified in *A.aegypti* and *D.melanogaster* are restricted to Diptera order (20 to 35% of all stage-specific genes), highlighting ongoing evolutionary processes that continue to shape life-stage transitions. Overall, our findings underscore the complex interplay between gene evolutionary age, expression specificity, and morphological transformations in development. These results suggest that the attraction of genes to critical life stage transitions is an ongoing process that may not be constant across evolutionary time or uniform between different lineages, offering new insights into the adaptability and diversification of dipteran genomes.

**Research Highlights:** Metazoan genomes contain instructions for a single cell to develop into a complete adult organism. We investigated stage-specific genes, i.e. genes that expressed only during certain developmental stages, in two dipteran species. We found that between 20 to 35% of stage-specific genes are active during transitions between major life stages and have emerged recently, suggesting an ongoing evolutionary process shaping life-stage transitions.

## Introduction

Metazoan genomes carry the complete set of instructions needed for a single cell to develop into a fully formed adult organism. However, not all genomic information is required or utilized at every stage of development. Instead, the progression through life stages is regulated by the dynamic expression of specific genomic regions, which ensures that molecular and biochemical functions are activated at the appropriate times to support the growth and development of the organism. Such precise regulation is particularly evident in holometabolous insects, whose evolutionary success is partly attributed to distinct lifestyles during juvenile and adult stages (Grimaldi & Engel, 2005; Marshall, 2012). These stages are not only morphologically distinct but also functionally specialized, enabling the insect to exploit different ecological niches throughout its life cycle. Drastic transformations between these stages are controlled by tightly regulated genome activity, which coordinates the timing and sequence of developmental processes.

The differential use of genomic information throughout development is well documented in the fruit fly *Drosophila melanogaster* (Arbeitman et al., 2002; Calderon et al., 2022; Holguera & Desplan, 2018; THE MODENCODE CONSORTIUM et al., 2010). Its genome encodes approximately 14,000 protein-coding genes, each exhibiting a distinct expression profile from the embryonic stage through to adulthood. While some genes, known as housekeeping genes, are expressed consistently across developmental stages due to their essential cellular functions, the majority of genes are selectively expressed in only a few specific developmental stages or tissues. These genes - referred to hereafter as stage-specific genes - include, for example, transcription factors and components of signaling pathways involved in processes such as organismal patterning, tissue specification, and cell differentiation, as well as genes with still unknown functions (Arbeitman et al., 2002). Interestingly, the temporal expression profiles of such stage-specific genes have been linked to their evolutionary age, with younger genes tending to display more restricted, stage-specific expression profiles (Heyn et al., 2014; Ma, Lau, & Zheng, 2024; Yang, Zou, Fu, & He, 2013).

Notably, genes that have originated recently often exhibit highly restricted expression patterns, frequently confined to specific tissues or particular stages of the life cycle (Brand & Ramírez, 2017; Gubala et al., 2017; Heames, Schmitz, & Bornberg-Bauer, 2020; Lange et al., 2021; Peng & Zhao, 2023; Zhou et al., 2015). This suggests that a transient and highly stage-specific gene expression profile during the time course of development may be a defining characteristic for evolutionarily young genes. However, it remains unclear whether the evolution of gene content and their associated expression profiles has followed a uniform pattern across different lineages and over evolutionary time. While studies have provided insights into genome evolution and gene expression dynamics of individual species (Akbari et al., 2013; Arbeitman et al., 2002; Gamez, Antoshechkin, Mendez-Sanchez, & Akbari, 2020; Gamez et al., 2020; THE MODENCODE CONSORTIUM et al., 2010), the extent to which these patterns are conserved or diverge across closely related species is still unclear.

Using the distinct, partitioned life stages of holometabolous insects as evolutionary context, we examined the distribution and evolutionary age of genes expressed transiently across developmental stages in two dipteran species, *Drosophila melanogaster* and *Aedes aegypti*. As flies, both species share typical traits of holometabolous development, i.e., transitions from embryo to larva, larva to pupa, and pupa to adult, prompting us to ask whether these transitions could be correlated with a shared set of new genes that originated in the stem group of holometabola and were specifically expressed during these conserved transitions. However, instead of identifying such signatures of shared innovations dating back to the stemgroup of holometabola, the comparison of these two distantly related fly species revealed much stronger signals of divergence within the insect order Diptera, highlighting independent gain of new genes with stage specific gene expression profiles.

## Results

### Life-stage transitions in *D.melanogaster* and *A.aegypti* are characterized by stage-specific gene expression

*D.melanogaster* is a well-studied holometabolous model system that features precisely defined transitions in its lifestyle (Bate & Arias, 1993). Developmental remodeling at the organismal and physiological level has been extensively documented (Campos-Ortega & Hartenstein, 1985). To test whether differential usage of its genome could be linked to particular stages of its lifestyle, we aimed to identify protein-coding genes with near-exclusive expression in specific time windows of development. To gauge such genome usage during the course of development in a way that would subsequently allow us to readily compare it with A. aegypti, we decided to use the expression of protein-coding genes as a first-level proxy.

To quantify the expression of protein-coding genes during specific time windows of *D.melanogaster* development, we used available modENCODE datasets (THE MODENCODE CONSORTIUM et al., 2010). The dataset comprises quantitative genome expression data for 24 hours of embryonic development sampled in consecutive two-hour intervals (12 embryonic stages), for each of the three larval stages, three stages of pupal development and 3 adult stages (THE MODENCODE CONSORTIUM et al., 2010). A comparable dataset is available for the mosquito *A. aegypti*; here the data comprises quantitative genome expression data for 76 hours of embryonic development, which has been sampled in consecutive four-hour intervals (20 embryonic stages), four larval stages, three stages of pupal development and 1 adult stage (Matthews et al. 2018). To confirm that the general timeline as well as individual windows of development could be reasonably compared between the two species as homologous ‘biological stages’, we examined the gene expression correlation of 3457 single-copy orthologous genes between the two species (Figure 1A, **Supplementary Table 3**). Based on these genes, notable correlations of gene expression could be detected throughout all stages of development, indicating the validity of reciprocal, homologous developmental stages in *D. melanogaster* and *A. aegypti*.

**Figure 1.**
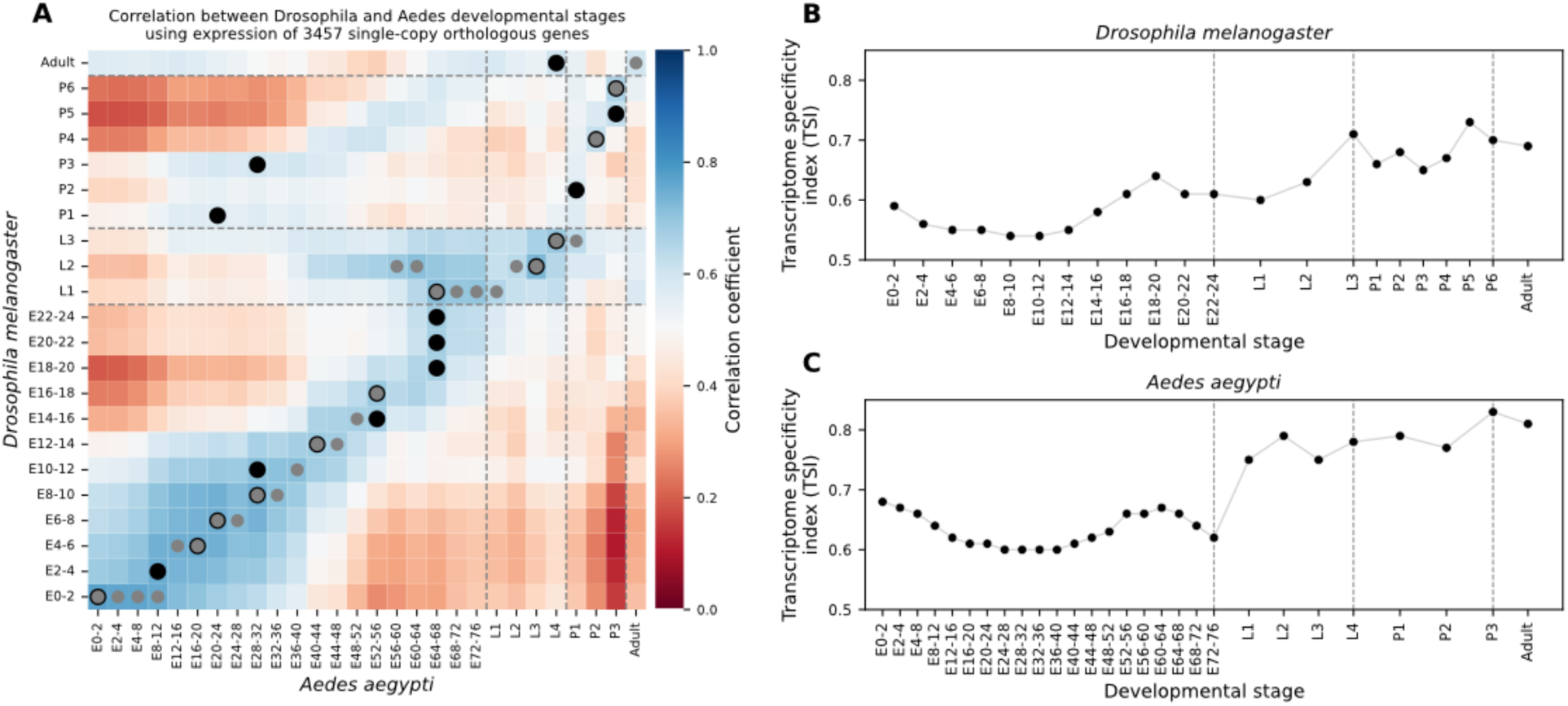
Comparative Analysis of Transcriptome Specificity in *D. melanogaster* and *A. aegypti*. (**A**) Gene expression correlation between *D.mealanogaster* and *A.aegypti*. The expression correlation was calculated by using 3,457 single-copy orthologues genes. The stage of maximal correlation in each species is marked (black circle: *D. melanogaster,* gray circle: *A. aegypti*). Expression profiles show notable cross-species correlation across developmental time, supporting the comparability of developmental stages between the two species. (**B, C**) Transcriptome specificity indices (TSIs) were calculated for each developmental stage in *D. melanogaster* (**B**) and *A. aegypti* (**C**). For each stage, the TSI reflects the summed tau-scores of all expressed genes, weighted by the proportion each gene contributes to total transcriptome output. TSIs are plotted over time across 19 stages in D. melanogaster and 22 stages in A. aegypti (E: embryo; L: larva; P: pupa).

To characterize differential gene expression in either species, previous work has successfully established the tau-scoring method, which was used to probe for differences in gene expression between tissues at one particular time interval (Yanai et al., 2005). To test whether the same methodology could also be applied to probe for differences in gene expression between different time points, we computed a tau-score for all protein-coding genes and their expression during the continuous developmental stages of *D.melanogaster* and *A.aegypti* (**Figure 1 Supplement 1, Supplementary Table 1**). We find that genes with a low tau-score are characterized by a broad distribution of gene expression that spreads over multiple developmental stages, while genes with a high tau-score show a very narrow distribution of expression, indicating temporal restriction. Taken together, our results suggest that - similar to the recently published approach by Lotharukpong et al. (Lotharukpong et al., 2024) - tau-scoring can be used to distinguish genes expressed in specific time intervals during the course of development. In the following we will refer to genes with high tau-scores (> 0.85) by the term “stage-specific” to distinguish them from genes with broader and temporally less specific expression patterns.

To assign each developmental stage of *D.melanogaster* with an overall estimate of gene expression specificity, we computed for each developmental stage a transcriptome specificity index (TSI), i.e., the normalized average tau score for all genes expressed at a given stage (Julien et al., 2012). When TSI was plotted as a function of developmental stages, it was found to change over time (**Figure 1A**): developmental episodes with higher TSI could be observed during early and late stages of embryonic development, as well as during larval and pupal development. Taken together, these results suggest that the transcriptome during the transition phases (embryo to larva, larva to pupa, pupa to adult) is characterized by more stage-specific gene expression than transcriptomes in the middle of life stages. This pattern is reminiscent of the hour-glass pattern that has been reported previously for the distribution of the average age of genes used during the course of development (Domazet-Lošo & Tautz, 2010).

To test the generality of trends observed in *D.melanogaster* in *Aedes aegypti*, we again used the expression of genes as proxy for genome usage and annotated each stage in this dataset by computing the TSI by using tau-scores. Like in *D.melanogaster*, TSI in *A.aegypti* varied during the course of development, indicating that the distribution of stage-specific gene usage was not distributed evenly (**Figure 1B**). Also like in *D.melanogaster*, early and late stages of *A.aegypti* embryonic development displayed high TSI values, while TSI values in mid-embryogenesis were low; during larval and pupal stages TSI and values were again high. These results for *A.aegypti* are thus comparable to those of *D.melanogaster* and support a general trend that transcriptome specificity appears to be higher at life-stage transitions. Taken together, our results suggest that life-stage transitions in flies - and possibly by extension in all holometabolous insects - are characterized by the expression of genes that are specific to life-stage transitions.

### Comparable life stages in *D.melanogaster* and *A.aegypti* share similar degrees of gene expression specificity

The notion that life-stage transitions in *D.melanogaster* and *A.aegypti* are enriched in the expression of stage-specific genes is based on the differential evaluation of TSI values across development in each of the two species. However, TSI is a composite measure based on the average tau-scores of hundreds, if not thousands of genes. Accordingly, TSI for any given developmental stage likely reflects a mixture of tau-scores from numerous genes with diverging specificity. To better understand the composition of signals contributing to the average transcriptomic tau-scores at various stages of development, we classified tau-scores of individual genes into different groups of increasing specificity. We then asked how much each group contributed to the global pattern of expression specificity during progressing stages of animal development.

When individual genes were analyzed for their expression profile and tau-score, genes with a tau-score value below 0.85 showed little if any enrichment of expression in specific stages, while genes above 0.85 were shown to be specific to developmental stages and yet could exhibit specific expression in more than one stage (**Figure 1 Supplement 1**). To further refine our classification of stage specificity, we thus interrogated three subpopulations of genes for their expression profiles across all developmental stages, i.e., genes with tau-scores of 0.85-0.9 (class I), 0.9-0.95 (class II), and 0.95-1 (class III). For *D.melanogaster* (**Figure 2 Supplement 1, A-C**), almost exclusive single-peak profiles were observed for class III genes (38%; 825 of 2496 genes with tau-score > 0.85), class II comprised genes with either one or two peaks (29%; 708/2496), and class I comprised genes that in many cases had more than two peaks in their developmental expression profiles (33%; 963/2496). For *A.aegypti* (**Figure 2 Supplement 1, D-F**), the overall pattern was similar: almost exclusively, single-peak profiles were observed for class III genes (30%; 1118 of 3118 genes with tau-score > 0.85), class II was comprised of genes with either one or two peaks (34%; 1053/3118), and class I comprised genes with more than two peaks (36%; 947/3118). Taken together, a tau-score value of 0.85 or higher suggests at least some stage specificity, both in *D.melanogaster* and *A.aegypti*.

**Figure 2:**
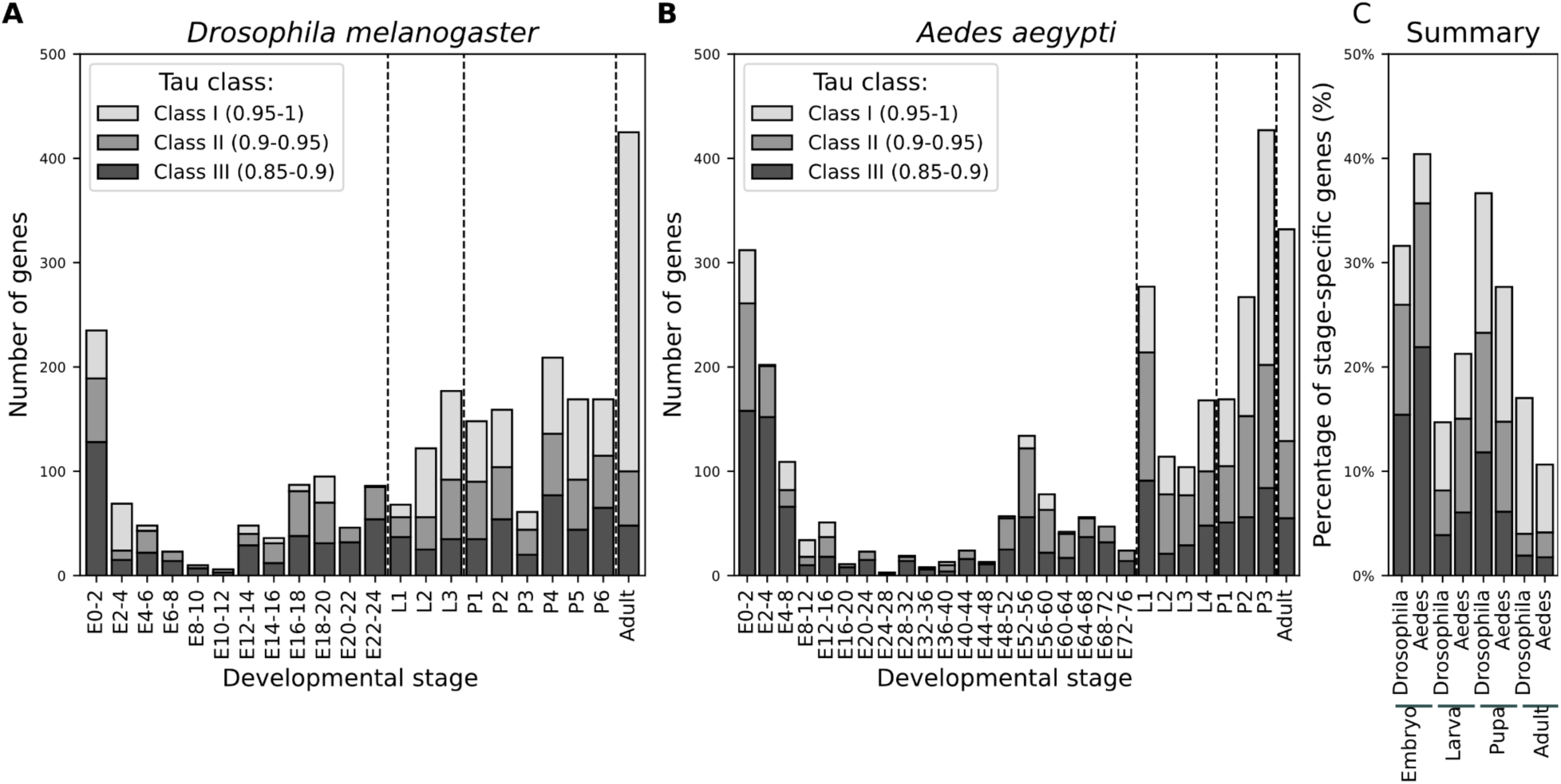
Number of genes with various levels of expression specificity in each developmental stage. (**A,B**) Expression of stage specific genes is shown as a composite of class I, class II, and class III genes for each stage of development in D.melanogaster (A) and A.aegypti (B). (**C**) The relative contribution of stage-specific genes from each class is shown for the four major life stages (embryo, larva, pupa or adult).

To address which of these gene subpopulations best explained the non-homogeneous TSI values over the course of development in *D.melanogaster* and *A.aegypti,* we selectively removed either class III genes, class II genes, or class I genes from the respective datasets. We then re-ran the TSI analysis and assessed how the overall profile changed (**Figure 2 Supplement 1**). For both species, we observed a notable decrease of TSI in the pupal stages after removal of class III genes (**Figure 2 Supplement 2**). During late embryogenesis and early larval stages, a substantial decrease of TSI could be observed after the removal of class II genes. Notably, the contribution of class I genes to the global TSI profile was low overall and in both species affected late embryogenesis and early larval stages.

To characterize the contribution of class I, class II, and class III genes to specific stages of development, we mapped their relative contributions to each developmental stage in both *D.melanogaster* and *A.aegypti* (**Figure 2**). In line with the global analysis of TSI (**Figure 2 Supplement 2**), we observed that, for both species, the relative contributions of class I, class II, and class III genes to individual stages were not uniform throughout the course of development. For example, some early embryonic stages in *D.melanogaster* (**Figure 2A, E2-4**) are characterized by the relative abundance of class III genes, whereas the last stages of embryogenesis show a high proportion of class I genes (**Figure 2A, E22-24**). Similar trends, albeit less pronounced, could be observed for *A.aegypti*. Using blastoderm formation as a known hallmark of embryonic development to register the different developmental speeds in *D.melanogaster* and *A.aegypti*, the same early embryonic stage in *A.aegypti* displayed an increased abundance of class III genes (**Figure 2B, E4-8**), while in later stages of embryonic development again showed a high proportion of class I genes (**Figure 2B, E68-76**).

While the total number of stage-specific genes was notably different between *D.melanogaster* and *A.aegypti* (2496 vs. 3118), the relative ratio of class I, class II, and class III across major life stages (embryo, larva, pupa, and adult) appeared relatively well conserved (**Figure 2C**). While we cannot formally exclude the possibility that such a pattern could stem from biases inherent to the temporal sampling of development (see discussion), we prefer to read these results as reflecting conservation of gene usage during early and late embryogenesis, separated by a mid-developmental transition (Levin et al., 2016). Collectively, our analyses indicate that regardless of variations in the number of stage-specific genes between the two species, the distribution of specifically expressed genes during major life stages is conserved in *D.melanogaster* and *A.aegypti*.

### High conservation and recent gain of stage-specific genes across dipteran phylogeny

Notably, the pupal stages in both species, *D.melanogaster* and *A.aegypti*, were characterized by the expression of genes that were exclusively active in this period of development (**Figure 2**). Pupal development is an exclusive feature of holometabolous development and absent in hemimetabolous insects (Grimaldi & Engel, 2005). Accordingly, the identified genes with pupal-specific expression in *D.melanogaster* and *A.aegypti* may be explained by at least two evolutionary scenarios. They could be ancient and present already in hemimetabolous genomes; in this case their exclusive expression observed during pupal development is a secondary innovation in *D.melanogaster* and *A.aegypti*. Alternatively, these genes emerged in the stem group of holometabola, be it *de novo* or via gene duplication, and potentially represent an innovation shared by all holometabola. To test these alternative evolutionary history scenarios, we dated the origin of stage-specific genes in *D.melanogaster* and *A.aegypti* using phylostratigraphy.

Phylostratigraphy dating of gene origin spanned over 450 million years of evolution and was based on 179 proteomes covering 6 phylostrata, including holometabola and arthropoda (**Figure 3**; **Supplementary Table 4 and 5**). With *D.melanogaster* as the focal group, we could date almost all stage-specific genes (2485 of 2496; the age of 11 genes remained ambiguous) and identified around 45% of these genes to be present in the stem group of insects (1141 of 2485 genes), i.e., prior to the innovation of holometabolous development. About a third of the dated stage-specific genes (904 of 2485 genes) originated within or at the base of Diptera, and only 3% (77 of 2485 genes) dated back to an origin in the stem group of holometabola (**Figure 3B**). When mapped onto individual time intervals of development, the proportions varied slightly, but the overall trend remained robust: for each developmental stage analyzed, on average, 70% of stage-specific genes that were found to be specifically expressed at a given stage of development originated outside of holometabola, while 18% originated within or at the base of Diptera (**Figure 3D**).

**Figure 3:**
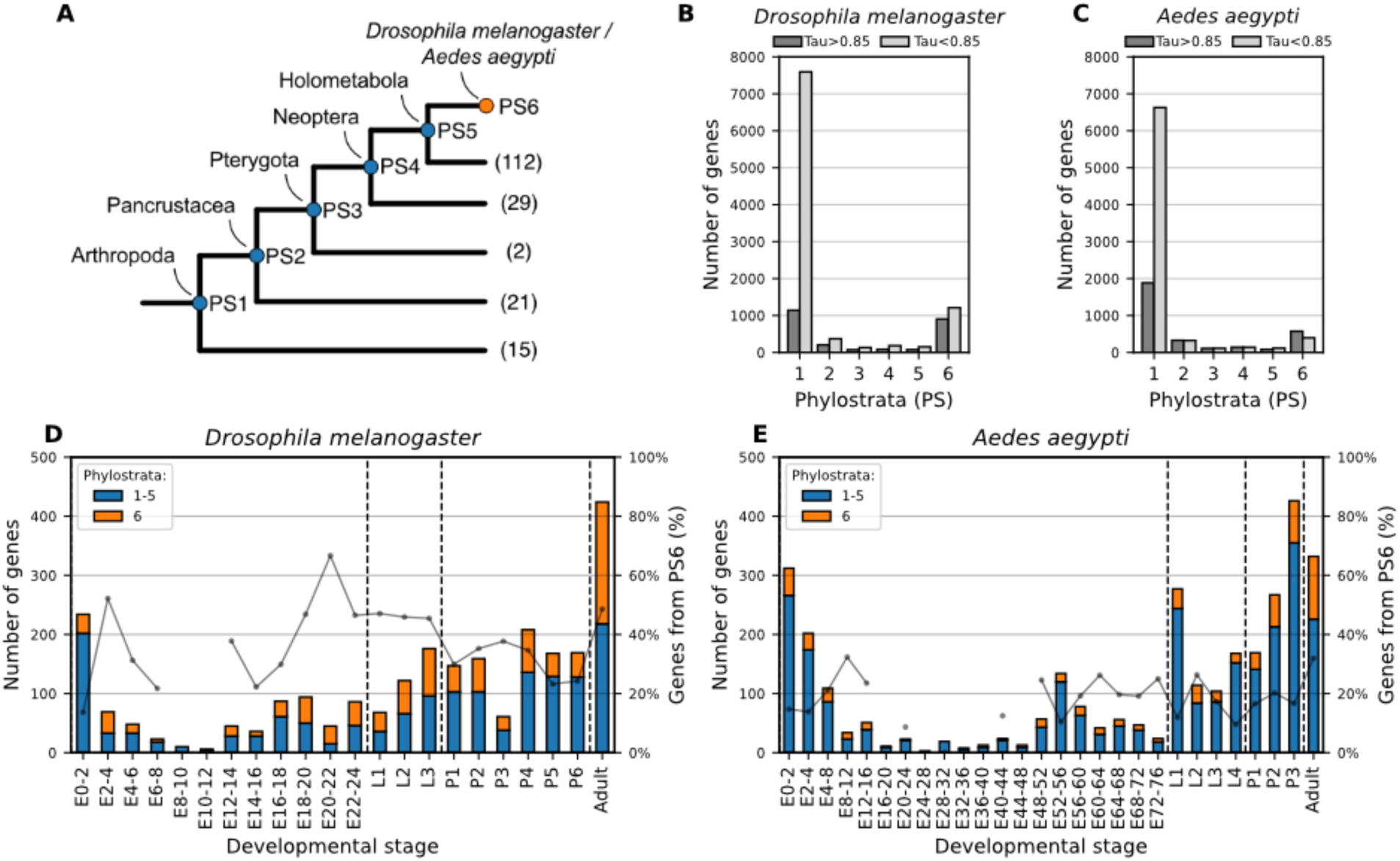
Phylostratigraphy analysis of stage-specific genes. (**A**) The evolutionary origin of stage-specific genes (class I to III) was mapped to 6 phylostrata in the insect phylogeny based on 179 proteomes across Arthropoda. The total number of proteomes supporting each phylostrata is shown in parenthesis. (**B-C**) Phylostratigraphic dating of stage-specific genes revealed a high proportion of genes originating in either phylostratum 1 (“Arthropoda-wide genes”) or phylostratum 6 (“diptera-specific genes”), with slightly different enrichment of age groups in *D.melanogaster* and A.aegypti (C). (**D, E**) All genes dating to either phylostrata 1-5 (blue) or phylostratum 6 (orange) were plotted as a function of developmental stage. The grey line indicates the ratio of diptera-specific genes (PS 6) to genes present outside Diptera (PS 1-5); to avoid spurious signal by noise, ratios for developmental stages with less than 30 genes were omitted.

Analogous phylostratigraphy analyses were performed using *A.aegypti* as the focal group. All but one stage-specific gene could be dated (3117 of 3118), and we identified around 70% of all these genes (1883 of 3117) as being present in the insecta stem group. About 20% of stage-specific genes could be dated back to an origin within or at the base of Diptera (573 of 3117 genes), and only 3% (83 of 3117 genes) dated back to an origin in the stem group of holometabola (**Figure 3C**). When mapped onto individual time intervals of development, the proportions varied slightly, but the overall trend remained robust: for each developmental stage analyzed, on average 77% of stage-specific genes expressed specifically at this stage originated outside of holometabola, while 23% originated within or at the base of Diptera (**Figure 3E**).

For both species, we tested whether the distribution of evolutionary gene age differs between stage-specific and stage non-specific genes using a chi-square test. This revealed a highly significant association between gene age and expression specificity in both species (D. melanogaster: χ² = 1092.66, df = 5, p < 2.2 × 10⁻¹⁶; A. aegypti: χ² = 890.74, df = 5, p < 2.2 × 10⁻¹⁶), indicating that stage-specific genes are significantly enriched for evolutionarily young genes. Unexpectedly, we find that many genes with highly specific expression profiles in *D.melanogaster* and *A.aegypti* have originated in the insect order of Diptera, either in the stem group or during its radiation, which is much more recent than, e.g., the transition from hemimetabola to holometabola.

### Rate of lineage-restricted gene gain differs across major Diptera families

The significant enrichment of evolutionarily young genes among stage-specific genes in both *D. melanogaster* and *A. aegypti* prompted us to ask when exactly these genes emerged within the insect phylogeny. To address this, we refined our phylostratigraphy analysis to increase temporal resolution within Diptera - the clade in which many of these genes appear to have originated. Specifically, we included 78 proteomes and 142 genomes covering 60 dipteran families (**Supplementary Table 4 and 6**). With *D. melanogaster* as focal group, this allowed us to resolve the origin of *D. melanogaster* stage specific genes with Dipteran origin (904 of 2485, see above) by dating them into 10 intra-dipteran phylostrata (**Figure 4A**). Notably, we found that only about 10% likely originated in the dipteran stem group about 250 million years ago, 10% in the stem group of Cyclorrhapha about 170 million years ago, 15% in the stem group of Schizophora at about 70 million years ago, while over 55% originated in the stem group of Drosophilidae or later (**Figure 4 A,B**). Taken together, these results suggest a notable gain of lineage-specific genes with high stage specificity since the divergence of Drosophilidae around 55-60 mya (Suvorov et al., 2022).

**Figure 4:**
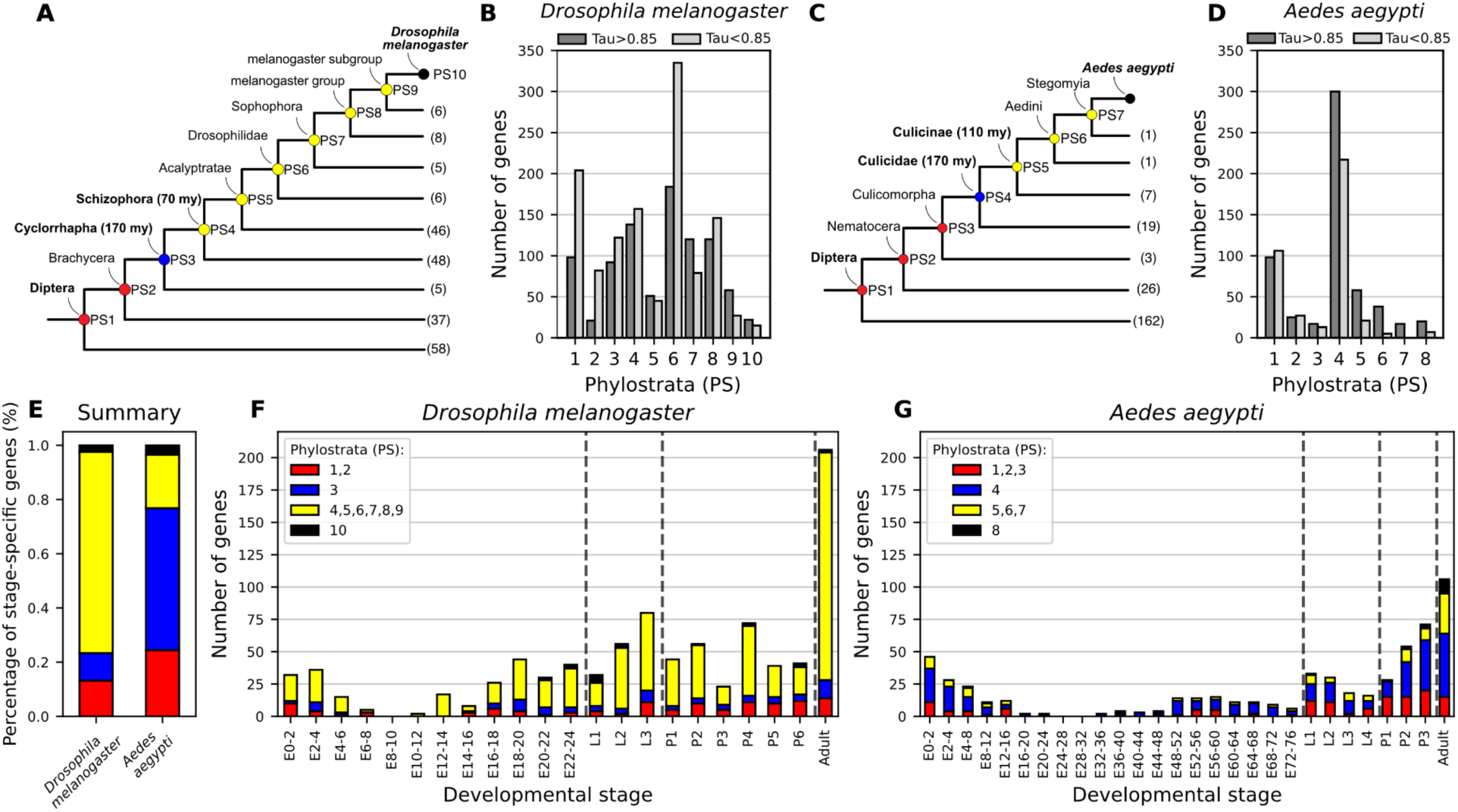
Stage-specific genes have been repeatedly gained in the recent evolutionary history of *D.melanogaster* and *A.aegypti*. (**A to D**) The evolutionary origin of Diptera-specific genes with stage-specific expression profiles was mapped to the dipteran phylogeny based on 93 proteomes and 146 genomes covering 60 taxonomic families. The genes were assigned to a total of 10 phylostrata for *D. melanogaster* (**A, B**) and 8 phylostrata for *A. aegypti* (**C**, **D**). For both species, the family-specific genes (Drosophilidae and Culicidae) mark the highest gain of stage-specific genes in both lineages. The total number of genomes and/or proteomes used to support each phylostrata is shown at the tips of the phylogenetic tree. (**E**) The distribution of stage-specific genes from age-binned phylostrata differs between *D.melanogaster* and *A.aegypt*i, with the *Drosophila* lineage showing a higher proportion of stage-specific gene gains in the last 70 million years compared to *Aedes*. (**F,G**) All the genes were plotted as a function of developmental stage for *Drosophila melanogaster* (**F**) or *Aedes aegypti* (**G**).

To resolve the origin of *A.aegypti* stage-specific genes with Dipteran origin (573 of 3118, see above), we first selected 8 phylostrata with comparable evolutionary age to the Cyclorrhapha/Schizophora/Drosophilidae lineage based on previously time calibrated phylogenies (da Silva et al., 2020). Specifically, we identified the stem group of Culicidae as rough age-equivalent for the stem group of Cyclorrhapha, and the stem group of Culicinae as age-equivalent for Schizophora (**Figure 4B**). By dating dating 557 of the 573 stage-specific genes with Dipteran origin (for 16 genes the age remained ambiguous) to the nodes in the in the *A.aegypti* lineage, we found that about 25% genes likely originated at the base of Diptera, 55% in the stem group of Culicidae about 170 million years ago, less than 1% in the stem group of Culicinae about 110 million years ago, while about 20% were species-specific and could not be resolved further for the past 110 million years (da Silva et al., 2020)(**Figure 4 A,C**). These results indicate that in *A.aegypti*, most of the stage-specific genes with Dipteran origin originated anytime after the emergence of the insect order and before the divergence of Culicidae, suggesting an age of roughly 170-250 million years.

When comparing stage-specific genes of dipteran origin in *D. melanogaster* and *Aedes aegypti*, we observed a notable difference in gene age distribution. Compared to *D. melanogaster*, the genome of *A. aegypti* contains proportionally more of the “older” stage-specific genes that can be traced back to the root of the Diptera. In contrast, *A. aegypti* harbors fewer genes that have emerged more recently - specifically, within the last 150 million years. This pattern suggests that *A. aegypti*, at least in terms of its stage-specific gene expression, may retain features of the ancestral dipteran genome more closely than *D. melanogaster* does. Vice versa, the distribution of stage-specific genes from different phylostrata indicates both a greater absolute number and a higher proportion of stage-specific gene gains in the last 70 million years in the Drosophila lineage compared to Aedes (**Figure 4 E-G**), suggesting more recent and extensive transcriptional innovation in Drosophila lineage.

### Stage-specific genes distribute evenly across the genome regardless of their age

The differences in the extent of genome evolution between *A.aegypti* and *D.melanogaster* may be, on the one hand, explained with overall diverging rates of evolution in the two lineages. Previous work has suggested rapid radiations in the stem group of Schizophoran flies about 65 million years ago (Wiegmann et al., 2011), which could explain the notable enrichment of stage- and lineage-specific genes that the *D.melanogaster* genome acquired in a period from about 70 to 4 million years ago. On the other hand, recent work has linked the rate of genome evolution with structural patterns of the genome (Benowitz et al., 2024), raising the possibility that the observed differences in the genome evolution between *A.aegypti* and *D.melanogaster* might be associated with genome structure.

To explore whether the genomes of *D.melanogaster* and *A.aegypti* - with regard to birth and expression of dipteran genes with a highly specific expression profile - could be associated with structural genomic differences, such as local clusters of stage-specific gene expression or hotspots of gene births, we explored whether the distribution of Dipteran-specific loci with stage-specific expression was randomly throughout the genome or appeared rather clustered in the same genomic region.

The genomes of *D.melanogaster* and *A.aegypti* differ in size by almost a factor of ten (*D.melanogaster*: 143MB; *A.aegypti*: 1.3GB). To ensure an even and comparable interrogation of genomic regions within and between the two species, we found the average size of topologically associated domains (TADs) to be a useful reference. TADs have been described as genomic domains that share a common neighborhood and often also regulation and similar expression. TADs scale with genome size and in *D.melanogaster* have an average size of about 150kb, while in *A.aegypti* TADs have an average size about 800 kb (Lukyanchikova et al., 2022; Szabo, Bantignies, & Cavalli, 2019). Based on TAD-sized windows, we subdivided the chromosomes of *D.melanogaster* and *A.aegypti* into non-overlapping windows over all chromosomes (*D.melanogaster*: 3 autosomes, 1 sex chromosome, 916 windows; *A.aegypti*: 3 autosomes, 1.2GB, 995 windows).

To detect local clusters of stage-specific gene expression in the genomes of the two species, we first assumed that stage-specific genes were distributed evenly among all genes in the genome. We then asked, for each genomic window, by how much the observed number of stage-specific genes in a given window deviated from the expected distribution (**Figure 5A, Supplementary Table 7**). In *D.melanogaster*, we detected <u>26</u> windows with a disproportional enrichment of stage-specific genes; in *A.aegypti*, <u>15</u> such windows were identified - both with up to fivefold more stage-specific genes than expected by chance (**Figure 5B and C**). This enrichment of genes with a stage-specific expression is notable and could be detected as highly significant statistical outliers in the genome of both species (**Figure 5AB**; **Figure 5 supplement 1**). On average, however, neither genome showed a global pattern for enrichment of genes with stage-specific expression (**Figure 5C**). Taken together, these results suggest that while stage-specific genes may occasionally cluster in specific genomic regions, the overall pattern does not reveal the presence of genome-wide structures that promote the accumulation of such genes in either species.

**Figure 5:**
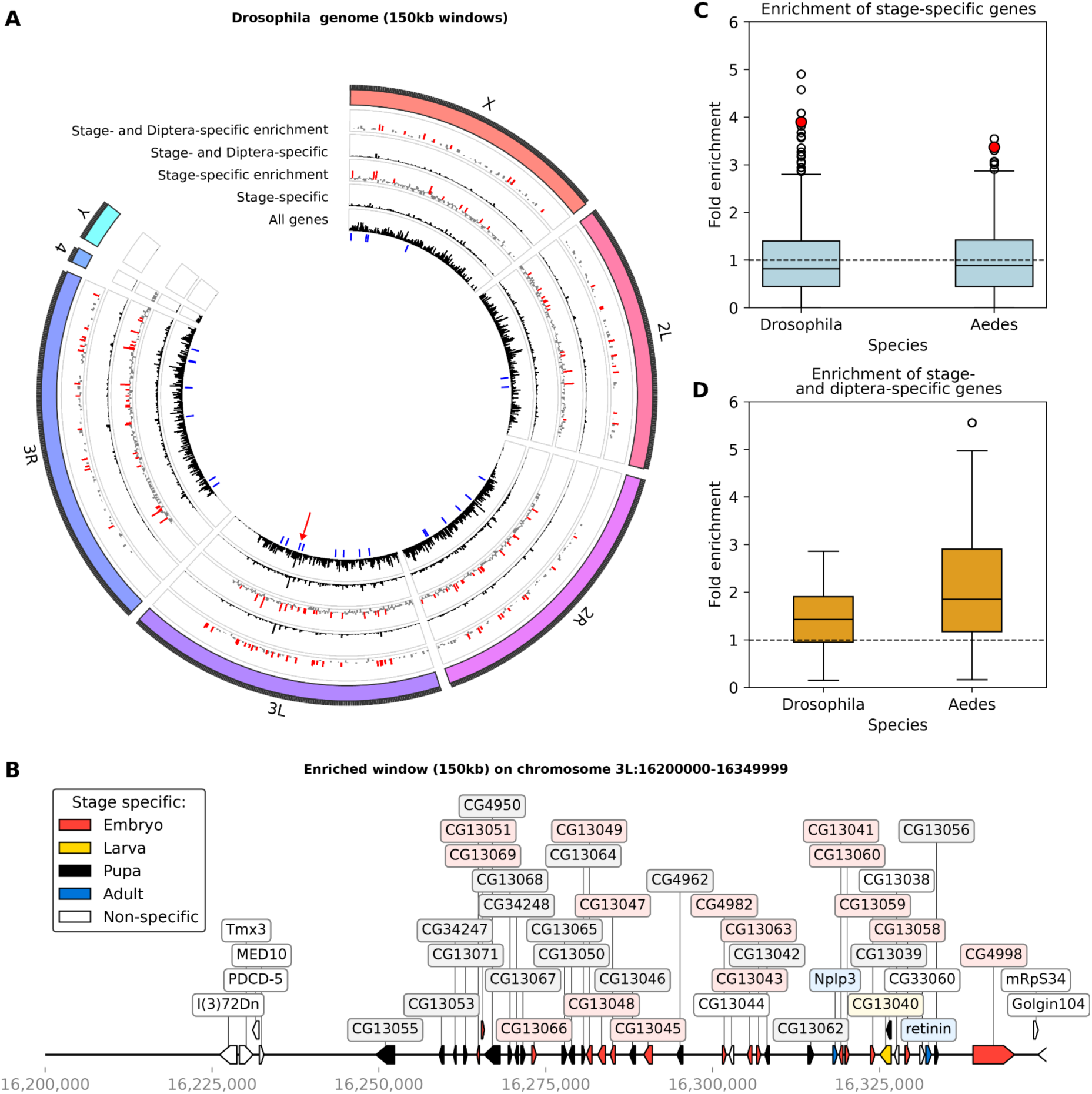
stage-specific genes can appear in genomics clusters. (**A**) Based on the average size of TAD, the chromosomes of *D.melanogaster* were divided into non-overlapping windows with size of 150kb (for *A.aegypti*, see Figure 5 Supplement 1). Shown for each genomic window are the total number of all genes (innermost ring, black), stage-specific genes (second inner ring, back), the relative enrichment of stage-specific genes (third ring, enrichment in red), the total number of stage-specific genes that originated within Diptera (fourth ring, black), and the relative enrichment of these genes within a given window (fifth ring, enrichment in red). (**B**) Example of a genomic locus with up to 5-fold enrichment of stage-specific genes from chromosome 3L illustrates the clustering of genes in a short genomic interval (the genomic position of the cluster is indicated as a red arrowhead in A; and as a filled red circle to indicate the outlier in C). (**C**) The average enrichment of stage-specific genes is comparable across genomic windows for *D.melanogaster* and *A.aegypti*; statistical outliers are marked as circles. (**D**) The average enrichment of stage-specific genes with dipteran origin across genomic windows for *D.melanogaster* and *A.aegypti* suggest that both species deviate from an expected random distribution; statistical outliers are marked as circles.

Recent population-level analyses in D. melanogaster have identified transposable element (TE)-rich genomic regions as hotspots for gene birth, with many recently emerged de novo genes overlapping regions of high TE density (Lebherz, Fouks, Schmidt, Bornberg-Bauer, & Grandchamp, 2024). To investigate whether stage-specific genes that originated within Diptera show a similar association, we compared the genomic distributions of TEs and dipteran stage-specific genes in the Drosophila genome. Using uniformly sized genomic windows, we quantified the relative enrichment of both features across the genome. While stage-specific genes showed a moderate correlation with the distribution of all protein-coding genes, no correlation was found with TE abundance (Figure 5 Supplement 2). These results suggest that, unlike young de novo genes, dipteran stage-specific genes are not preferentially located in TE-rich regions.

To detect possible hotspots of gene birth among the Dipteran genes with stage-specific expression more generally, we assumed that every gene with stage-specific expression had the same probability of having emerged in the dipteran stem group or later. We then asked, for each genomic window, by how much the actual observed number of newly emerged genes deviated from this expected distribution (**Figure 5A**, **Figure 5 supplement 1, Supplementary Table 7**). For *D.melanogaster*, we did not detect any genomic window in which stage-specific genes were significantly enriched for a recent birth in the dipteran stem group or later. For *A.aegypti*, by contrast, we identified 19 genomic windows that were significantly enriched for stage-specific genes of recent origin (**Figure 5D**).

Taken together, our results suggest that hotspots of gene births may exist among the subclass of genes with stage-specific expression profiles, at least in the genome of A.aegypti. While further analyses will be required to understand the evolutionary mechanisms leading to such clusters, they appear overall to be extremely rare and are thus unlikely to constitute a structural element that could explain diverging rates of genome evolution between D.melanogaster and A.aegypti.

## Discussion

Selective utilization of genomic information across different life stages is a defining feature of metazoan organisms, most notably exemplified during developmental processes (Arbeitman et al., 2002; THE MODENCODE CONSORTIUM et al., 2010). To start to understand how such selective genome usage changes and adapts during evolution, we have used the expression of protein coding genes as a first-level proxy to compare genome usage over the course of development in two different fly species, *Drosophila melanogaster* and *Aedes aegypti*. As flies, both species are characterized by shared transitions in life stages from embryo to larva, from larva to pupa, and from pupa to adult, raising the possibility to uncover shared characteristics of genome usage in both flies. Surprisingly, however, rather than identifying signatures of shared ancestral commonalities, we find much stronger signals in recent divergence of selective genome usage.

### Lineage restricted innovations in fly genomes appear to be expressed at life stage transitions

While a comprehensive analysis based on a subset of approximately 4,000 genes has previously highlighted the diversity of *D.melanogaster* genome utilization over the course of development (Arbeitman et al., 2002), less attention has been given to possible signatures of underlying genome evolution. To begin to address this gap, we compared genome utilization over the course of development using the full gene complement of *D.melanogaster* (consisting of about 14,000 protein-coding genes sampled over 22 life stages), and *A.aegypti* (consisting of about 14,500 protein-coding genes sampled over 27 life stages). To assess expression specificity across developmental stages, we used tau-scoring, a computational methodology that was originally developed to evaluate tissue-specific gene expression. As has been recently demonstrated also for other, independently and more recently evolved multicellular organisms (Lotharukpong et al., 2024), we could confirm the utility of tau-scoring for the detection of genes with stage-specific expression. Because the approach showed limitations in identifying genes with multiple expression peaks and restricted the effectiveness in capturing complex temporal patterns, we used a cumulative transcriptome specificity index as described recently (Lotharukpong et al., 2024). Based on this cumulative scoring approach, we could show that transcriptome specificity was notably higher during life-stage transitions compared to mid-stage periods, both in *D.melanogaster* and *A.aegypti*.

When plotted against developmental time in *D. melanogaster* and *A. aegypti*, both transcriptome specificity and the age distribution of stage-specific genes peak at the transitions between major life stages and dip in the intervening periods - remarkably recapitulating a classical “hourglass” profile (**Figures 1B,C; 3D,E; 4F,G**). This hourglass pattern - high divergence in early and late stages but strong conservation during the phylotypic period - has been observed for multicellular taxa at all scales, between strains, in populations, between species, and even across kingdoms, for both gene-expression specificity and transcriptomic age (Domazet-Lošo & Tautz, 2010; Duboule, 1994; Kalinka et al., 2010; Levin, Hashimshony, Wagner, & Yanai, 2012a; Liu & Robinson-Rechavi, 2020; Lotharukpong et al., 2024; Quint et al., 2012; Zalts & Yanai, 2017). Mechanistically, strong purifying selection at the phylotypic stage is often invoked to maintain a core “bauplan” (Zalts & Yanai, 2017). However, several studies also report accelerated evolution (positive selection) in early and late development, for both protein sequences (Coronado-Zamora, Salvador-Martínez, Castellano, Barbadilla, & Salazar-Ciudad, 2019; Liu & Robinson-Rechavi, 2018b, 2018a) and enhancers (Liu et al., 2021). Over deep phylogenetic distances, however, high sequence turnover presumably erodes such signals, making positive selection increasingly difficult to detect with standard sequence-based methods. In this context, the repeated recruitment of newly originated genes into specific developmental stages - particularly outside the phylotypic stage - may offer an additional, and potentially more sensitive, proxy for identifying episodes of adaptive diversification. While many new genes likely arise neutrally or under weak selection, their consistent expression in early or late stages across divergent lineages suggests that these life-stage transitions may provide permissive and selectively favorable contexts for gene integration. In this view, stage transitions function as “innovation sandboxes,” where novel genes are tested, refined, and, if advantageous, retained and then potentially redeployed elsewhere in development. Under this model, the hourglass profile of new, stage-specific genes could signal ongoing adaptive diversification at evolutionary distances beyond the reach of classical sequence tests. However, alternative explanations would need to be considered, such as mechanistic biases in gene emergence or chromatin accessibility, as they could influence where new genes appear. Future work combining functional assays of young genes and high-resolution chromatin maps will be essential to disentangle neutral drift from positive selection in shaping the evolving developmental genome.

### Ancient life stage transitions continue to attract new genes at variable rates

The transition from hemi- to holometabolous development dates back about 350 million years and has been considered an ancient evolutionary innovation (Grimaldi & Engel, 2005; Misof et al., 2014). If a life stage like the pupa, which is specific to holometabolous development and does not occur in hemimetabolous insects, is found to be enriched for lineage restricted innovations in the genomes of *D.melanogaster* and *A.aegypti*, it may be tempting to imagine how new genes that are specifically expressed during this stage are of a similar age and functionally related to its innovation. Contrary to such expectations, however, many of the genes that we identified as new and specific during the pupal stages, in the genome of *D.melanogaster*, originated much more recently than the transition from hemi- to holometabolous development.

To explain this discrepancy, several possibilities can be considered. First, even though stage-specific genes are highly significantly enriched for “young” genes that originated within the insect order of Diptera, we cannot exclude the possibility that the finding could be an artifact of methodological limitations. The apparent enrichment of Dipteran-specific genes might reflect biases in genome annotation quality across species. Dipterans, and Drosophila in particular, are extensively studied and often serve as reference genomes; as a result, other insect genomes may be under-annotated or annotated with a Drosophila-centric bias. This could artificially inflate the number of genes appearing to originate in Diptera. Second, a more biological explanation relates to the divergent rates of genome evolution in different insect lineages. Dipteran genomes, particularly those of Drosophila, may be inherently more dynamic - undergoing higher rates of gene birth and turnover. This genomic fluidity could contribute to the accumulation of lineage-specific, stage-specific genes, independent of ancient developmental innovations. Third, the signal of gene age and stage-specificity may simply erode over time. Genes that initially emerged with stage-specific expression - perhaps in association with the origin of novel life stages like the pupa - may have been gradually co-opted by other stages of development. As these genes became integrated into broader regulatory networks, their specificity may have faded, rendering them undetectable in our analysis. Taken together, we favor a biological explanation in which the enrichment of genes at particular stages is not necessarily a reflection of the stage’s evolutionary age. Instead, certain stages may act as recurring points of gene recruitment throughout evolution, continuing to attract new, stage-specific genes long after their initial emergence.

The analysis of *D.melanogaster* and *A.aegypti* genome utilization for specific developmental life stages suggests that the attraction of new genes may not be constant across evolution or equal between different branches of the dipteran phylogeny: about 70 to 4 million years ago, the *D.melanogaster* genome accumulated much more new stage-specific genes in a period in which the *A.aegypti* genome experienced little to no change at all. These differences in the accumulation of new genes with stage-specific expression could not be associated with obvious structural differences, suggesting that they likely result from known episodic radiations in the dipteran lineage. Such radiation occurred in particular at the base of Diptera, in the subgroups of Brachycera and Schizophora (Wiegmann et al., 2011). Each of these radiations coincided with the appearance of novel phenotypic features, such as long proboscis for nectar feeding in Brachycera and a novel escape mechanism from puparium in Schizophora (Wiegmann et al., 2011). Since its divergence from the last common ancestor of Diptera, the genome of *D.melanogaster* has been shaped by these radiations, the *A.aegypti* genome has not. Taken together, our results suggest that - in comparison with *D.melanogaster* - *A.aegypti* represents a genome that much more closely resembles the ancestral dipteran genome.

### Limitations of the study

This study is, at least in part, based on a definition of developmental stages by temporal windows. However, binning a continuous developmental process into distinct temporal windows is inherently prone to artifacts; it is based on time intervals that do not necessarily reflect biological stages (Levin, Hashimshony, Wagner, & Yanai, 2012b). Because beginning and end of ‘true’ biological stages remain unknown and can - at least in the current study - only be approximated by non-overlapping temporal developmental episodes of two hours each, we cannot exclude the possibility that some aspects of the reported stage-specificity of gene expressions result from an inherent structural bias in the used datasets. Structural bias in the used datasets may stem from two principle errors that come with an approximation of developmental episodes by temporal windows of limited sampling and resolution: a biological stage may be temporally undersampled if a two-hour window of development in fact comprises the equivalent of two or more biological stages, or it may be temporally oversampled if a given biological stage takes longer than the two-hour sample interval, such that the process is artificially segregated into two or more two-hour time windows. As discussed below, temporal undersampling and oversampling have different consequences.

In case of temporal undersampling, two or more ‘true’ developmental stages, which each may have very specific but functionally and developmentally unrelated gene expression profiles, are getting accidentally binned into one temporal window. Using tau-scoring under such conditions may artificially inflate the number of genes picked up with high stage-specificity because one time interval represents two or more biological stages. Notably, this type of bias is not limited to fast developmental processes that are temporally undersampled. A similar bias may also be expected if a particular time interval comprises multiple independent but co-occurring biological processes, e.g., during the completion of various organ formation programs in the final stages of embryonic development. Consequently, it may be possible to observe artificially inflated numbers of stage-specific genes under conditions in which multiple stages are aggregated into one time interval. Critical intervals to look into may be, e.g., late embryogenesis and final organ differentiation. In case of temporal oversampling, the principle error leads to the temporal dispersal of a single extended developmental process into two or more two-hour intervals. Using tau-scoring under such conditions will artificially deflate the number of genes that will be specifically associated with this developmental process: even though genes may be specific and exclusive to such an extended developmental process, they may be considered non-specific because they have peaks in three consecutive stages. Consequently, it may be possible to observe artificially deflated numbers of stage-specific genes under conditions in which a biological process is temporally oversampled. Critical intervals to look into may be, e.g. the phylotypic stage or mid-embryogenesis in general.

Despite these potential limitations, the core findings of our study remain robust. Although temporal undersampling and oversampling could introduce artifacts in comprehensively identifying every stage-specific gene, the broader patterns we observed - such as the principal enrichment of lineage-specific innovations during life-stage transitions and the divergence in genome usage between *D.melanogaster* and *A.aegypti* were supported consistently. While further refinement of temporal resolution could enhance precision, the fundamental insights into genome evolution are not compromised by these methodological constraints.

## Conclusions

In conclusion, our comparative analysis of genome utilization across developmental stages in *D.melanogaster* and *A.aegypti* reveals important insights into the evolutionary dynamics of gene expression. While we initially anticipated uncovering shared ancestral patterns of genome usage associated with holometabolous development, our findings instead highlight the prominent role of recent, lineage-specific innovations. The enrichment of new, stage-specific genes at life-stage transitions suggests that these transitions continue to attract new genetic elements even long after their initial evolutionary emergence. Together, these results underscore the complex interplay between evolutionary age, gene expression specificity, and the ongoing adaptation of genomes to developmental demands, providing a clearer picture of how genome evolution shapes the unique life histories of metazoan species.

## Methods

### Life cycle transcriptomes

Publicly available and pre-processed transcriptomes covering different life stages were downloaded for *D.melanogaster* (THE MODENCODE CONSORTIUM et al., 2010) and *A.aegypti* (Matthews et al., 2018). We downloaded processed transcriptomes for which the sequencing reads had been mapped and the gene counts inferred in RPKM (Reads Per Kilobase Million). The Drosophila transcriptome was based on FlyBase annotation release 6 assembly and the *A.aegytpi* transcriptome was based on GCF_002204515.2_AaegL5.0 assembly. The *A.aegypti* transcriptome was combined by the previous authors from multiple sources, including (Matthews et al., 2018) and (Akbari et al., 2013) and mapped against the GCF_002204515.2_AaegL5.0 assembly. Samples covering relevant developmental life-stages (embryo, larva, pupa and adult development) were extracted from the datasets. In total we collected 22 RNA-seq samples covering all four life stages of development of *D.melanogaster* and 27 RNA-seq samples covering three life stages of development of A.aegypti (**Supplementary Table 1 and 2**). Due to the absence of an RNA-seq sample of the whole adult in the *A.aegypti* dataset, we used the gene expression of adult carcass (the tissue of the adult body without legs and head) as an equivalent of the *A.aegypti* adult stage. Aedes carcass expression data originates from the same dataset as all the other stages. Furthermore, to allow for comparison between *A.aegypti* and *D.melanogaster* life cycle transcriptomes, RNA-seq samples of different adult life-stages of D.melanogaster, that is RNA-seq samples of 1-, 5- and 30-day old adult flies, were merged into a single sample by taking average expression.

Life-cycle transcriptomes were further processed by merging sample replicates, filtering out non protein-coding genes and low expressed genes. For both species, RNAseq reads from identical life-stages were merged by computing an average expression level. Then, to allow for intra- and inter-species comparison of life-stage transcriptomes, the expression levels in both datasets were converted to transcript per million (TPM). Next, all the protein-coding genes with below significant expression (TPM < 10) across all life-stage were filtered out from the dataset. After data processing, the *D.melanogaster* life cycle transcriptome dataset consisted of 22 life-stages and 12157 protein-coding genes; the *A.aegypti* life cycle transcriptome dataset was composed of 28 life-stages and 10845 protein-coding genes (**Supplementary Table 3**).

### Gene expression correlation analysis

To determine the best matching reciprocal life-stages between *Drosophila melanogaster* and *Aedes aegypti* we used gene expression correlation analysis of single-copy orthologous genes. To determine single-copy orthologous genes between *D.melanogaster* and *A.aegypti*, we used Orthofinder v2.5.5 with default parameters (Emms & Kelly, 2019). For Orthofinder input we used longest isoform proteomes of both species. Orthofinder detected a total of 3457 single-copy orthologous genes between the two species (**Supplementary Table 4**). Gene expression correlation was calculated based on Pearson product-moment correlation coefficient using previously TPM normalized gene expression values calculated in this study.

### Transcriptome specificity index

To characterize transcriptome specificity index (TSI), we computed the stage-specificity for each protein-coding gene in life-stage specific transcriptomes of *D.melanogaster* and *A.aegypti* using tau-scores (Julien et al., 2012; Yanai et al., 2005)(**Supplementary Table 3**):

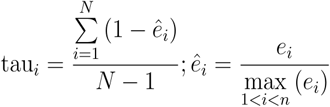

Here, *n* represents all the stages of the life cycle, *i* represents all the genes used. Tau-score value ranges from 0 to 1; 0 marks low and 1 high gene expression stage-specificity.

To characterize how broadly or transiently genes were expressed in each developmental stages, we calculated transcriptome specificity index (TSI) for each life-stage in both species (Lotharukpong et al., 2024). TSI was calculated as a tau-score normalized relative to the average gene expression for each developmental stage:

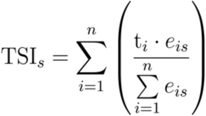

Here, *s* represents a developmental stage, *n* represents all the genes expressed during developmental stage, t represents tau value of a gene and i represents all the genes expressed in one developmental stage.

### Phylostratigraphy

We inferred the evolutionary age of stage-specific protein-coding genes for *D. melanogaster* and *A. aegypti* using genetic phylostratigraphy with Genera v1.4.2 software (Barrera-Redondo, Lotharukpong, Drost, & Coelho, 2023). Genera performs sequence homology searches of input protein sequences against specified proteomes or genomes (in our case selected Arthropoda species proteomes and genomes). Based on species’ phylogenetic relationships and gene presence-absence patterns, each gene is assigned to the most distant phylostratum (PS) in which it can be detected. Genes showing spurious presence-absence patterns are labeled as “contamination” and excluded from analysis.

We conducted two independent phylostratigraphy analyses. First, we determined gene ages across Arthropoda using 179 proteomes and 6 phylostrata (PS 1 representing Diptera-specific genes; PS6 representing Arthropoda-specific genes; **Supplementary Table 3 and 5**). For genes with multiple isoforms, we used the longest isoform. After analyzing all protein-coding genes with above-threshold expression (TPM>10), we successfully assigned ages to 12,156 Drosophila and 10,844 Aedes protein-coding genes. Of these, 2,116 Drosophila genes and 969 Aedes genes were classified as Diptera-specific.

Second, we date the age of Diptera-specific with higher resolution in Diptera phylogeny by using additional 78 proteomes and 142 genomes covering 60 Diptera families (**Supplementary Table 3 and 6**). Based on available genomic and proteomic data, we could assign the Diptera-specific genes either into 10 or 8 phylostrata for Drosophila or Aedes, respectively. For genes with multiple isoforms, we used the longest isoform.

### Genome cluster analysis

To detect genomic clustering of stage-specific protein-coding genes we organized genomic elements into non-overlapping windows and calculated fold-enrichment of genetic elements per window. Genome assemblies and annotation files for D. melanogaster (GCF_000001215.4) and A. aegypti (GCF_002204515.2) were downloaded from NCBI. Chromosomes were divided into 150 kb windows for D. melanogaster (916 windows) and 800 kb windows for A. aegypti (995 windows). Each protein-coding gene was assigned to a single genomic window based on its position; genes spanning two windows were allocated to the window containing most of the gene’s length. Additionally, for Drosophila, we determined the location of transposable elements from the assembly annotation file and assigned them into genomic windows.

Enrichment of stage-specific genes was calculated using a random distribution model, where we multiplied a global representation value (ratio of stage-specific to all genes) by the total gene count per window (**Supplementary Table 7 and 8**). Fold-enrichment was determined by normalizing the stage-specific gene count to this window-specific representation value. A similar approach was applied to calculate enrichment of transposable elements and stage- and Diptera-specific genes, assuming random distribution among stage-specific genes. Total gene counts and enrichment values were mapped to the genome using pyCircos v0.3.0, and highly enriched windows were visualized with DnaFeaturesViewer v3.1.3.

### Data visualization

Data was visualized with custom python, R scripts and Affinity Designer. All bar plots and line plots were made by using matplotlib v3.8.4 or seaborn v0.13.2. Heatmaps were made using seaborn v0.13.2.

## Acknowledgements

The authors acknowledge support by the state of Baden-Württemberg through bwHPC and funding by the Deutsche Forschungsgemeinschaft (LE 2787/4-1 and STA 850/60-1) as part of SPP 2349 (GEVOL).

---

**Figure 1 Supplement 1:**
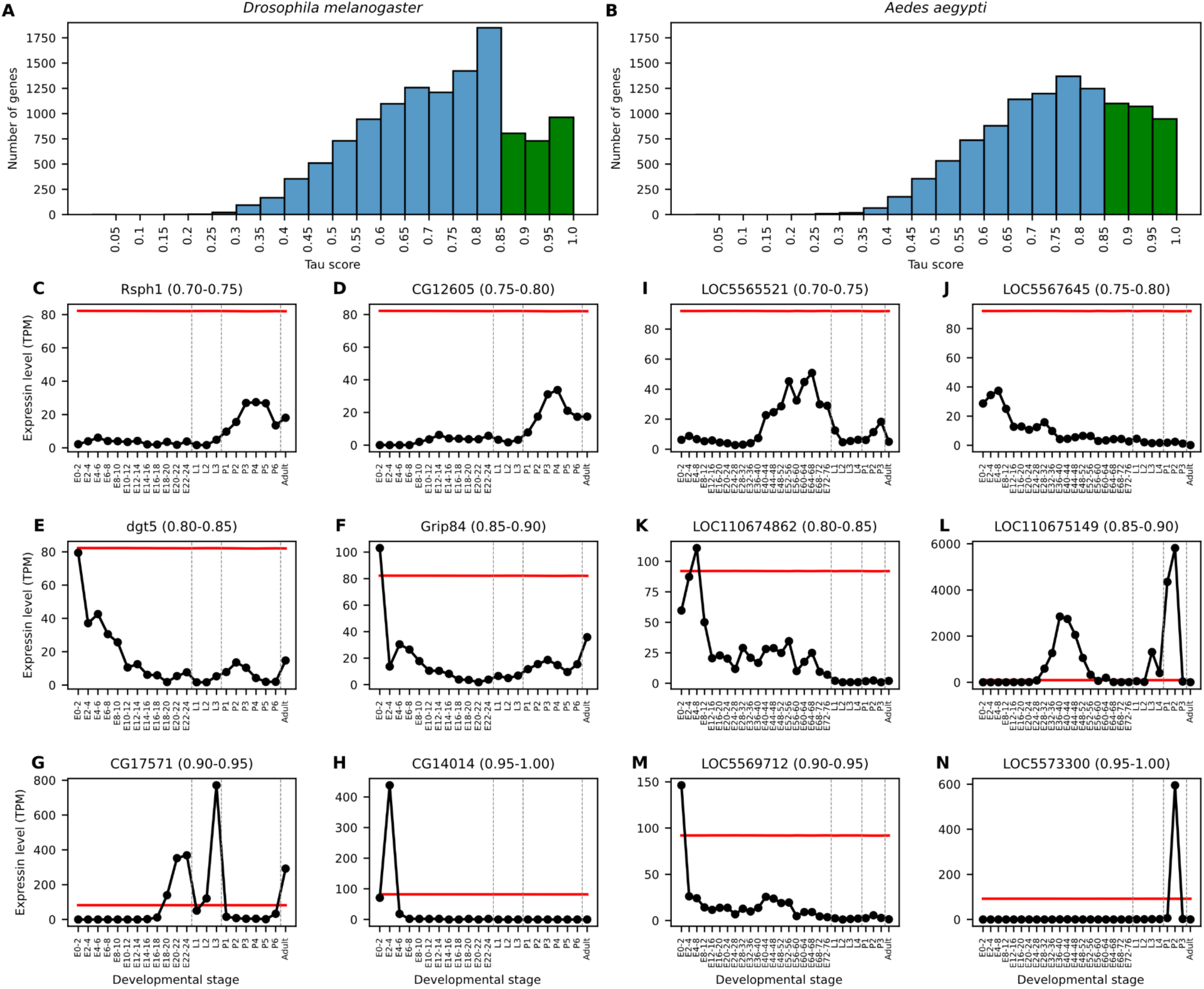
Classification of stage-specific genes according to tau-score values. (**A-B**) To score for how strongly transient expression levels were increased during specific stages of development, a tau-score was computed for each gene over the course of development in *D.melanogaster* (**A**) and *A.aegypti* (**B**). Tau scores were based on developmental transcriptomic data that comprised 20 stages in *D.melanogaster*, and 25 in *A.aegypti*. Based on their tau-scores, genes were grouped into 20 bins of 0.05 intervals from 0.00-0.05 to 0.95-1.00. (**C-N**) To illustrate the expression profiles of individual genes comprised in the top six tau-score bins (0.70-0.75, C,I; 0.75-0.80, D,J; 0.80-0.85, E,K; 0.85-0.90, F,L; 0.90-0.95, G,M; 0.95-1.00, H,N), for each bin the temporal expression profile of a randomly picked gene is plotted for *D.melanogaster* (**C** to **H**) and *A.aegypti* (**I** to **N**). FlyBase gene symbols (*D.melanogaster*) and NCBI IDs (*A.aegypti*) and the respective tau-score intervals are indicated on top of each panel. Gene expression levels are normalized as transcript per million (TPM); the red lines mark the mean TPMs computed for all genes expressed in each developmental stage.

**Figure 2 Supplement 1:**
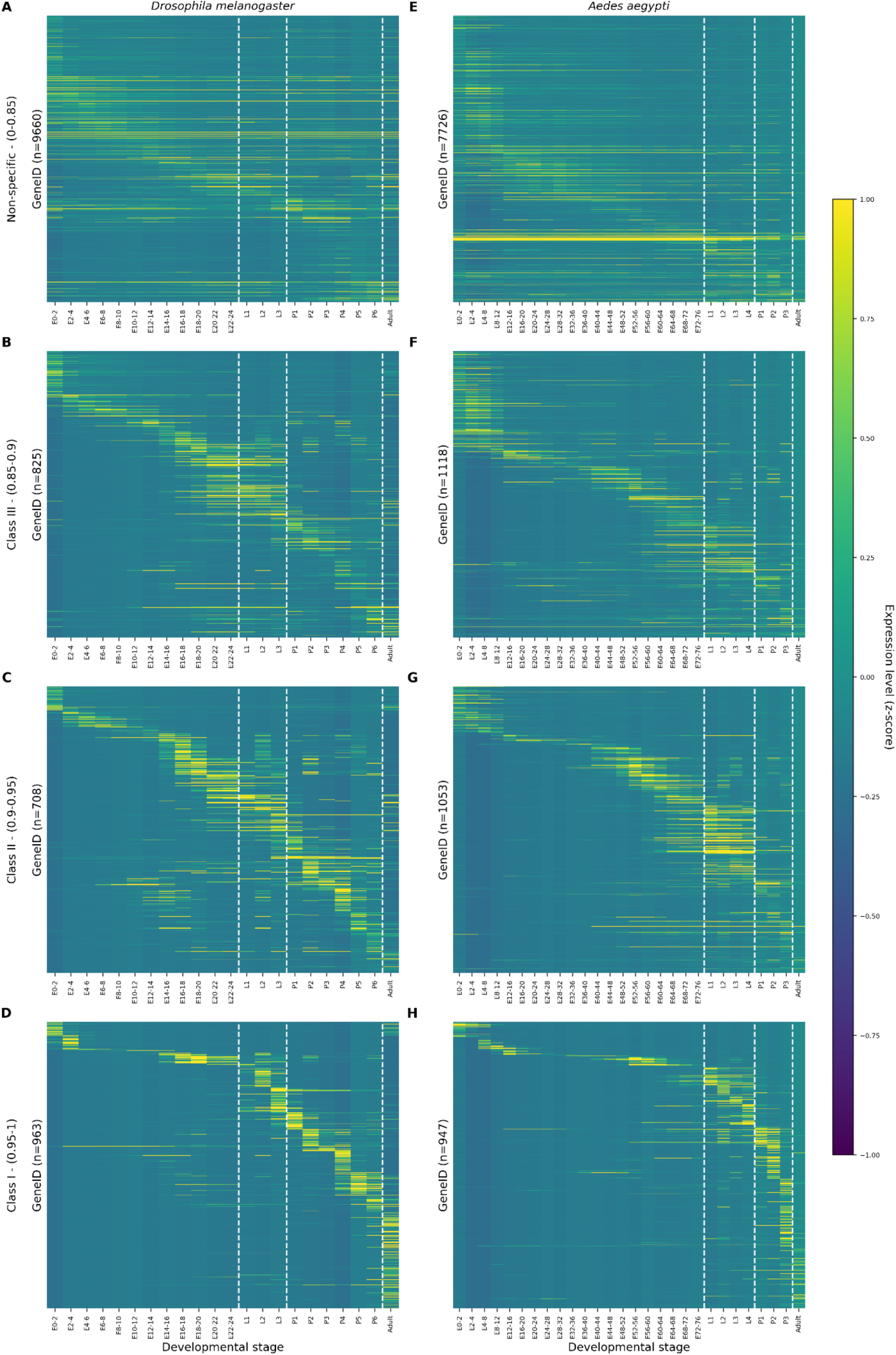
Tau-scoring effectively distinguishes genes with single or multiple peaks of expression across developmental stages. (A-H) All protein-coding genes from *D.melanogaster* (A-D) and *A.aegypti* (D-F) were grouped into four classes of increasing stage specificity (non-specific, 0-0.85; class I, 0.85-0.9; class II, 0.9-0.95; class III, 0.95-1). Most class III genes are expressed transiently in a single developmental stage (D and H), class II genes are mostly expressed in one or two developmental stages (C, G), and class I genes are expressed in up to five developmental stages (B, F). Expression levels for each gene are indicated as standard deviation from its mean expression over the time course of development (z-scores) using heatmap color coding.

**Figure 2 supplement 2:**
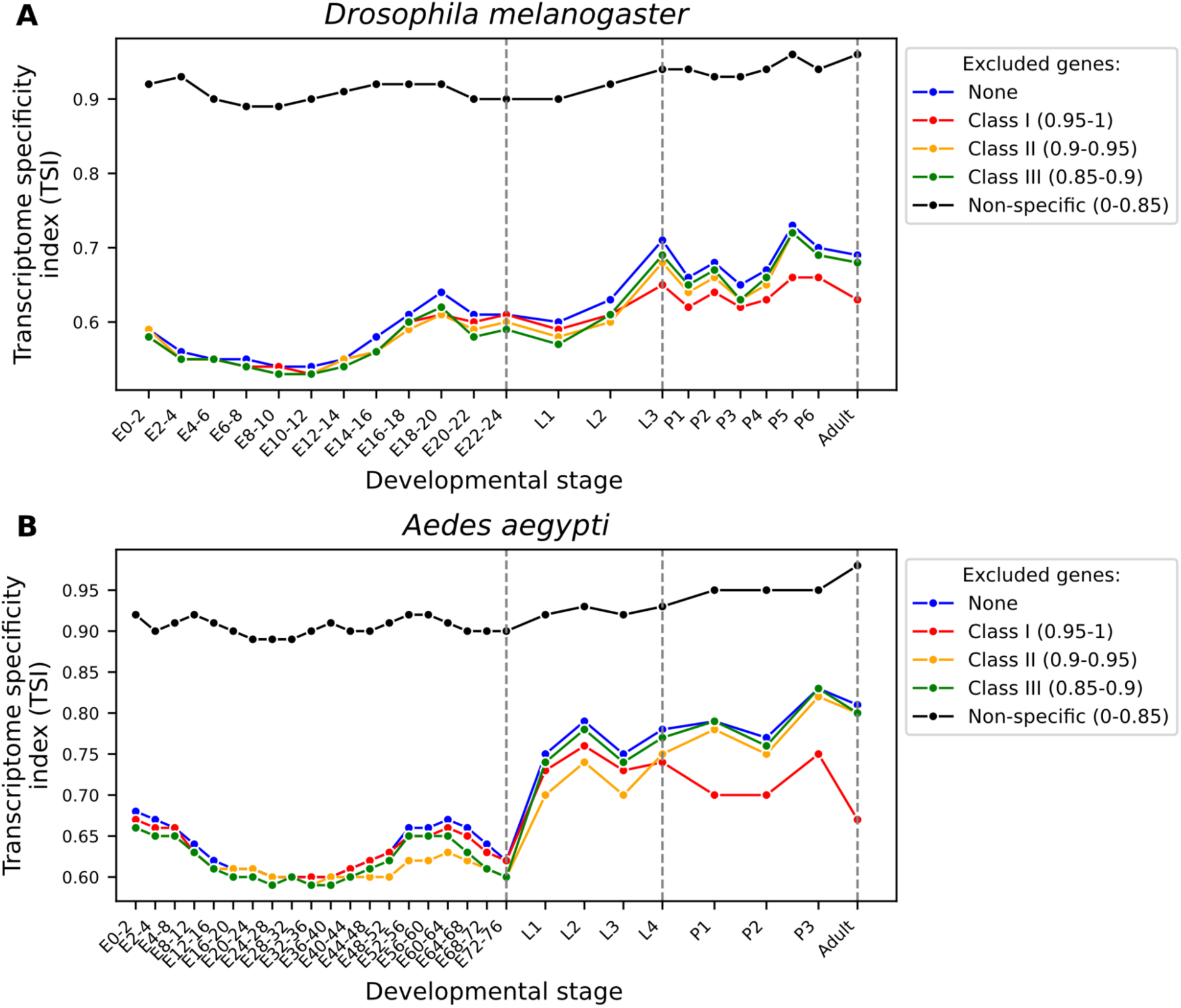
TSI profiles change when classes of genes with different expression specificity are selectively excluded from the analysis. (**A,B**) TSI scores were plotted over the course of development in *D.melanogaster* (A) and *A.aegypt*i (B), either without any gene class removed (“None”), after removal of class III genes (0.95-1; red), class II genes (0.90-0.95; yellow), class I genes (0.85-0.90; green) or after removal of all non-specifically expressed genes (“Non-specific”, black).

**Figure 5 Supplement 1:**
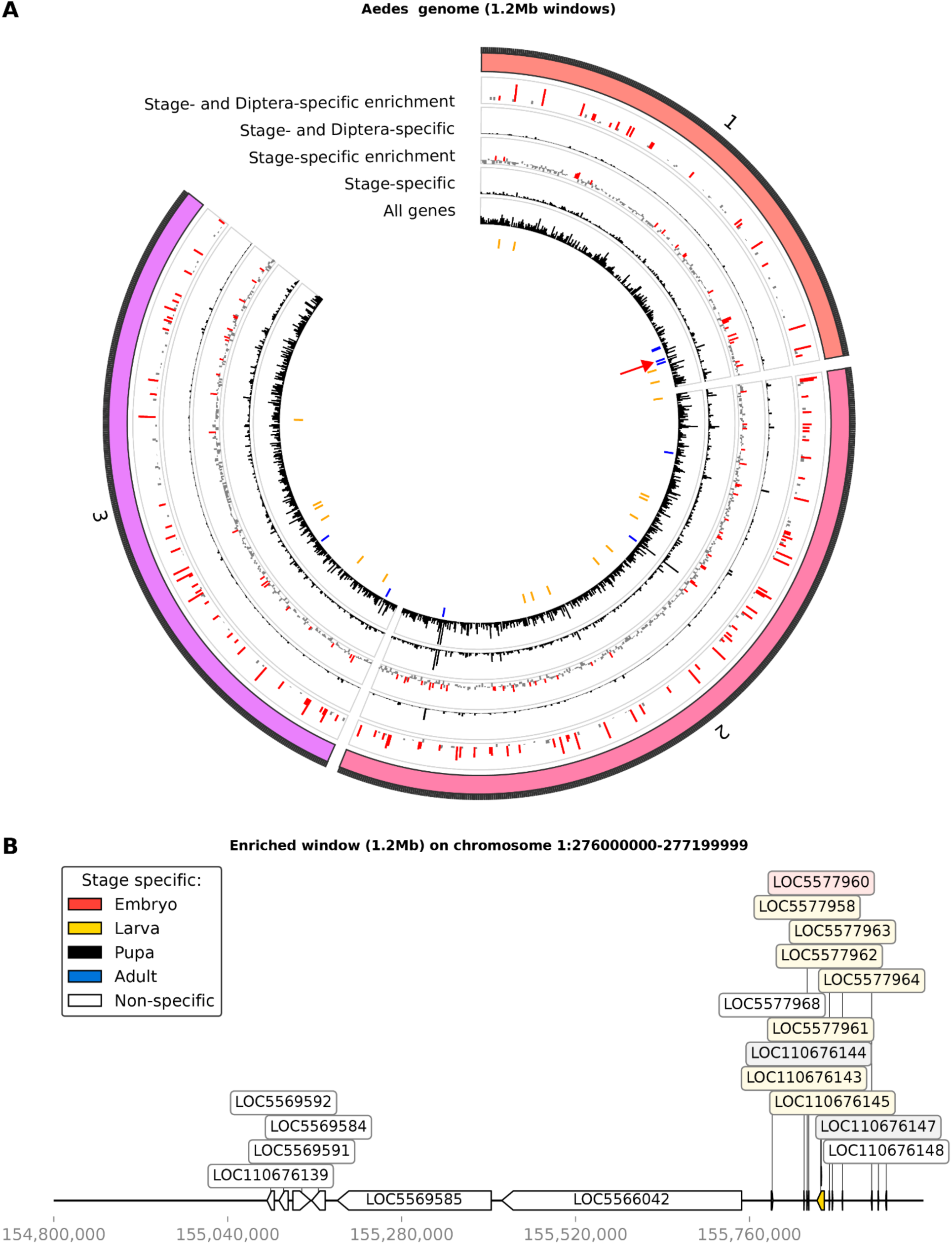
Clusters of stage-specific genes in the genome of *A.aegypti*. (**A**) Based on the average size of TAD, the chromosomes of *A.aegypti* were divided into non-overlapping windows with size of 800kb. Shown for each genomic window are the total number of all genes (innermost ring, black), stage-specific genes (second inner ring, black), the relative enrichment of stage-specific genes (third ring, enrichment in red), the total number of stage-specific genes that originated within Diptera (fourth ring, black), and the relative enrichment of these genes within a given window (fifth ring, enrichment in red). (**B**) Example of a genomic locus with up to 5-fold enrichment of stage-specific genes from chromosome 1 illustrates the clustering of genes in a short genomic interval (the genomic position of the cluster is indicated as a red arrowhead in A; and in Figure 5C as a filled red circle to indicate the outlier).

**Figure 5 Supplement 2:**
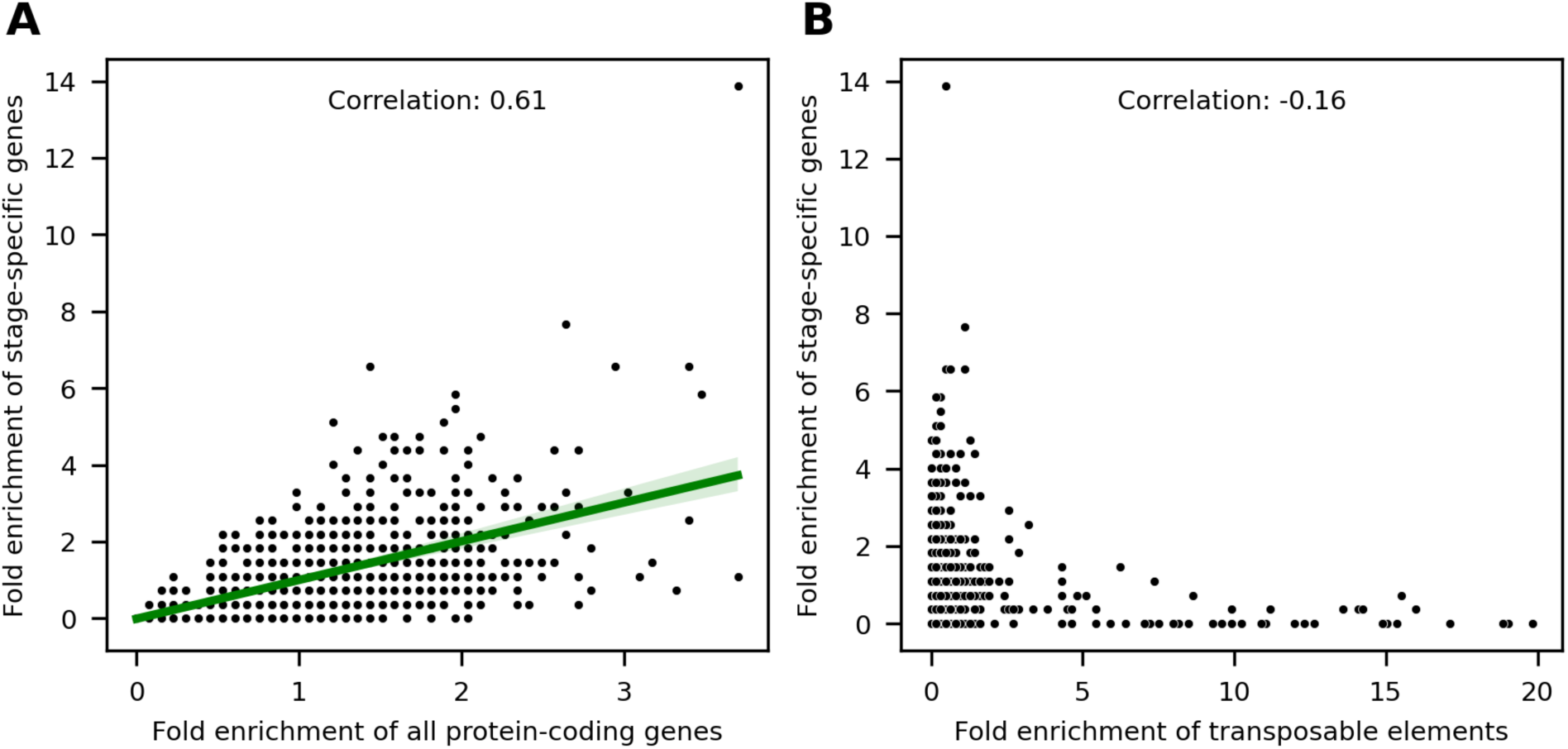
Enrichment of stage-specific genes in D.melanogaster genomic windows shows moderate correlation with protein-coding genes, but not with transposable elements. (**A**) The correlation between relative enrichment of protein-coding genes (x-axis) and stage-specific genes (y-axis) across Drosophila genomic windows (In total of 917 genomic windows). The enrichment of stage-specific genes is in moderate correlation with the enrichment of protein-coding genes (Correlation coefficient 0.61). (**B**) The correlation between relative enrichment of transposable elements (x-axis) and stage-specific genes (y-axis) across Drosophila genomic windows. The enrichment of stage-specific genes is not correlating with transposable elements (Correlation coefficient −0.16).

